# Variability in the diet diversity of catfish highlighted through DNA barcoding

**DOI:** 10.1101/2020.09.18.268888

**Authors:** Chinnamani Prasannakumar, Gunasekaran Iyyapparajanarasimapallavan, M. Ashiq Ur Rahman, P. Mohanchander, T. Sudhakar, K. Kadharsha, K. Feroz Khan, J. Vijaylaxmi, Narra Prasanthi, Kumaran Subramanian, Seerangan Manokaran

## Abstract

Identification and quantification of fish diet diversity was the first step in understanding the food web dynamics and ecosystem energetics, where the contribution of DNA barcoding technique has been important. We used DNA barcoding to identify the stomach contents of a euryhaline, benthophagous catfish *Ariius maculatus*. From 40 catfish stomach items sampled in two different seasons, we barcoded 67 piscine and macro-invertebrates prey items, identified as belonging to 13 species in 4 major phyla (viz., Chordate, Arthropod, Annelida and Mollusca). It is important to note that the mollusc taxa *(Meritrix meritrix* and *Perna viridis)* and a species of fish *(Stolephorus indicus)* could not be found among the gut contents of *A. maculatus* sampled during the pre- and post-monsoon season, respectively. Among the piscine diets of *A. maculatus, Eubleekeria splendens* (23.5%) and *Stolephorus indicus* (23.5%) were the major prey taxa during pre-monsoon season. The hermit crabs forms the major constituents of both pre- and post-monsoon seasons, among invertebrate taxa. Polychaete, *Capitella capitata* (25.92%) was abundantly consumed invertebrates next to hermit crabs. We noticed that in pre-monsoon *A. maculatus* was more piscivorous than post-monsoon. As revealed through Kimura-2 parametric pairwise distance analysis, the diet diversity was relatively higher in post-monsoon. The accumulation curve estimated 57 haplotypes within 14 barcoded species (including the host *A. maculatus)*. Majority of haplotypes were found among fishes (47.36%) followed by Arthropods (28.07%), Annelids (14.03%) and Mollusca (10.52%), respectively. This study also highlights that there is a growing concern about *A. maculatus’s* aggressive predation on commercially important stocks of fish and invertebrates. We will continue to expand the coverage of species barcoded in the reference database, which will become more significant as meta- and environmental DNA barcoding techniques become cheaper and prevalent.

## 1. Introduction

In Ichthyology, the first step towards identifying trophic levels of the fish species was the determination and quantification of fish stomach contents (Hynes, 1950; Hyslop, 1980; Cresson et al., 2014) and understanding of food web dynamics and energy of the ecosystem (Beauchamp et al. 2007). The traditional visual survey of stomach content, however, did not provide data to be compared across studies (Cortes, 1997; Hernandez, 2019), as level of taxonomic resolution at which a prey was classified (e.g. order, family, genus or species level) and the metrics used for quantification (e.g. count, volume) differed between the studies (Berg, 1979; Hyslop, 1980; Hansson, 1998; Hernandez, 2019). While much work has been expended in standardizing dietary analysis (Pinkas, 1971; Cortes, 1997), it has not been universally accepted as a standard technique (Baker et al., 2014). Prey items that is present at different digestive stages when analysing the gut produces ambiguity in taxonomic resolution and identification. These findings differ through studies and rely on different factors, such as the methodology adapted for prey identification, prey condition, prior taxonomic knowledge of the prey species, and the objective of the gut analysis (example: Elliott, 1967; Baker and Sheaves, 2005; Saunders et al., 2012; Gray et al., 2015). In addition to complications due to unreported prey assumption, variable taxonomic resolutions compound the inconsistency and make the findings incomparable between studies.

DNA barcoding refers to sequencing the standard barcode region and matching them with archived sequences derived from validated species to facilitate species identification (Hebert et al., 2003; Joly et al., 2014; Kress et al., 2015). DNA barcoding has been used to study trophic dynamics (Valdez-Moreno et al. 2012; Wirta et al. 2014; Moran et al. 2015), environmental forensics (Dalton and Kotze 2011; Handy et al. 2011; Gonsalves et al. 2015), cryptic diversity and invasive species identification (Hebert et al. 2004; Conway et al. 2014; Bariche et al. 2015), ecosystem and evolutionary diversity evaluation (Ward et al. 2005; Baldwin et al. 2011; Weigt et al. 2012a; Leray and Knowlton 2015) and phylogenetic exploration (Nagy et al. 2012; Baeza and Fuentes 2013; Betancur-Ret al. 2013). While a number of markers available (e.g., 16sRNA, 18sRNA, matK, rbcL, ITS), a ∼ 650 base pair (bp) region in the mitochondrial cytochrome c oxidase 1 (COI) gene is one of the most widely used in fish (and other animals) (Ward et al., 2005; Lakra et al., 2010; Khan et al., 2011; Weigt et al. 2012b). Species recognition is made by comparing the query sequence to that of archived sequences in reference databases, such as the Barcode of Life Database (BOLD) (Ratnasingham and Hebert 2007) using the BOLD-Identification System (BOLD-IDs) and in GenBank (National Center for Biotechnology Information) using Basic Local Alignment Sequence Tool (BLAST) (Altschul et al. 1990). DNA barcoding has been widely applied in assessment of diversity and composition of fin and shell fishes (Teodoro et al., 2016; Sharawy et al., 2017; Zhang et al., 2017; Wang et al., 2017; Kuguru et al., 2018; Ran et al., 2020; Xu et al., 2021). DNA barcoding is currently commonly used in consumer markets to identify sharks and their products (Holmes et al., 2009; Nijman et al., 2015; Hellberg et al., 2019). DNA barcodes acts as a potential forensic tool to track illegally traded and mislabeled endangered fishery products (Pappalardo & Ferrito, 2015; Di Pinto et al., 2015; Sembiring et al., 2015; Velez-Zuazo et al., 2015; Carvalho et al., 2017; Bunholi et al., 2018).

In recent years, the use of DNA barcoding to identify digested prey items has increased, especially in the identification of deepwater sharks prey items (Barnett et al. 2010; Dunn et al. 2010); Lake fish predators (Carreon-Martinez et al. 2011); invasive lionfish *Pterois* sp. (Valdez-Moreno et al. 2012; Cote et al. 2013; Rocha et al. 2015; Dahl et al., 2017; Ritger et al., 2020; Santamaria et al., 2020); groundfish (Paquin et al. 2014); pterygophagous (fin eating; Arroyave and Stiassny 2014) and lepidophagous (scale eating; Boileau et al. 2015) fishes; introduced largemouth Bass *Micropterus salmoides* (Jo et al. 2014); warm-water catfish (Moran et al. 2015); herbivorous juvenile Sandy Spinefoot *Siganus fuscescens* (Chelsky Budarf et al. 2011); gray seals *Halichoerus grypus* and harbor porpoises *Phocoena phocoena* (Méheust et al. 2015) and stranded Humboldt squid *Dosidicus gigas* (Braid et al. 2012). Nonetheless, few studies have investigated the effectiveness of DNA barcoding in relation to the digestive state of fish prey (Carreon-Martinez et al. 2011; Moran et al. 2015). Although few studies have successfully used DNA barcoding in the analysis of catfish piscine prey items of (Moran et al., 2015; Aguillar et al., 2016; Guillerault et al., 2017), studies are rare in elucidating both vertebrate and invertebrate prey content of benthophagous catfishes like *Arius maculatus* (Thunberg 1792). A. *maculatus* is a euryhaline, benthophagous species in tropical and sub-tropical waters, estuaries, rivers and coastal regions (Mazlan et al., 2008; Chu et al., 2011; Jumawan et al., 2020), whose economic importance and potential for aquaculture have recently been recognised (Jumawan et al., 2020). Commonly referred as spotted catfish (Chu et al., 2011), *A. maculatus* belongs to Ariidae family (Carpenter and Niem 1999), along with eight other species and is known for its pharmaceutical and nutraceutical values (Al-Bow et al., 1997; Osman et al., 2007). The present study aims to identify the species composition in the dietary items of *A. maculatus* occurring in the Vellar estuary (southeast coast of India).

The effectiveness of DNA barcoding for species identification largely depends on the establishment of broad and robust barcode reference databases of validated, verified species with vouchered specimens. A lack of vouchered or incorrectly identified sequences (Ekrem et al. 2007; Valdez-Moreno et al. 2012; Weigt et al. 2012a) would severely decrease the utility of reference databases. In addition, to capture potential genetic variation, including undocumented cryptic diversity, it is important to sequence an adequate number of individuals from across a species range (Weigt et al. 2012b). We have made considerable efforts in the past decade as part of the Indian Census of Marine Life (ICoML) to recover barcodes in reasonable numbers of marine phyla including fin & shell fishes, invertebrates (Khan et al., 2010, 2011; PrasannaKumar et al., 2012; Thirumaraiselvi et al., 2015; Rajthilak et al., 2015; Rahman et al., 2013, 2019; Hemalatha et al., 2016; Palanisamy et al., 2020; PrasannaKumar et al., 2020a, b; Manikantan et al., 2020; Thangaraj et al., 2020) and plants (Sahu et al., 2016; Narra et al., 2020) occurring in and around the Vellar estuary. Hence we predict a high rate of success in identification of dietary items of *A. maculatus* occurring in this environment.

## 2. Material and methods

### 2.1. Catfish collection and stomach analysis

During April (pre-monsoon) and December (post-monsoon) 2011, catfish sampling was carried out in the Vellar estuary (latitude: 11° 29’N; longitude: 79° 46’E). Individuals were collected fresh from local fishery folks who used hand nets to fish regularly in Vellar estuary. A total of 40 catfishes were collected (18 in pre-monsoon; 22 in post-monsoon). Upon collection freshly captured fish were transported to DNA barcoding Lab, Centre of Advanced Study in Marine Biology, Annamalai University, within 1 km from collection site, in the saltwater ice slurry to slow down catfish digestion and prey DNA degradation (Baker et al., 2014). Each catfish has been measured (to nearest mm; total length (Lt)) and weighed (to nearest 0.1g).

After examination of the oesophagus and gills for prey, Catfish *(Arius maculatus* (Thunberg, 1792)) digestive tracts were removed. In order to remove digestive enzymes, excess chyme and particulate organic matter using 500 μm sieves, the digestive tracks were dissected and their contents were rinsed with RO water. Recognisable prey taxa were divided into two classes, namely fish and invertebrates, and into 2 digestive stages, namely whole animals (i.e., most of the biomass was present) and digestive remnants (in bits and pieces). The remnants were rinsed once in 100% molecular grade ethanol (Sigma) and approximately 3 mm^3^ tissue plug was exercised under the ∼1 mm top tissue layer (especially in fish preys) from all recognisable prey items (examined under binocular microscope whenever necessary) to avoid sample contamination by *A. maculatus* cells (Cote et al., 2013). Once again, the tissues exercised were rinsed and stored in microfuge tubes (Thermo Fisher Scientific) containing 100% ethanol at 4°C until further analysis. Residual tissues in scalpel and forceps were removed with 95% ethanol and flame sterilized between each exercise. Samples of lateral tissue samples were also taken from *A. maculatus* for DNA barcoding because if cannibalism was alleged in the diet, it would provide useful reference (Valdez-Moreno et al., 2012).

### 2.2. DNA extraction, PCR and sequencing the dietary contents

The DNA was extracted using the DNeasy Blood & Tissue Kits (Qiagen), following the manufacturer’s instructions or standard protocols (Prasannakumar et al., 2020). Tissue was placed in an extraction buffer containing proteinase K and homogenized with polypropylene disposable sterile pestles (Thermo Fisher Scientific). The homogeneous was digested at 56 °C until complete digestion of the sample (i.e, when most of the homogenate is more translucent) that was within 12 hours. Elution buffer was used as the negative control to test the purity of the reagents. The extracted DNA for quantification and purity estimation (i.e., using 260/280 nm ratio) was quantified using Nanodrop (Thermo Fisher Scientific). Elution buffer provided in the kit was used during Nanodrop measurements to calibrate the blank. Also, DNA concentration was checked in 1.2% Agarose gels and the high DNA yields (>85 ng/μl) were 10X diluted before PCR in ultrapure water.

DNA from fish-like prey samples was PCR amplified using COI primers; FishF1 and FishR1 (Ward et al., 2005) and invertebrate samples were using LCO1490 and HCO2198 (Folmer et al., 1994). A reaction mixture volume of 25μl was used to conduct PCR; 12.5μl Taq PCR Master Mix (Invitrogen, India), 11μl distilled water, 0.5μl forward primer (10 μl, 0.5μl reverse primer (10 μM), and 0.5μl DNA template (50-80 ng/μl) were used. Initial steps of 2 min at 95 °C, followed by 35 cycles of 30 s at 94 °C, 30 s at 54 °C, and 1 min at 72 °C, followed by 10 min at 72 °C and kept at 4 °C were used for FishF1/FishR1 primer amplified samples. Initial denaturation for 2 min at 95 °C, followed by 5 cycles at 94 °C for 30 s, 46 °C for 45 s, 72 °C for 45 s and 35 cycles at 94 °C for 30 s, 51°C for 45 s, 72 °C for 45 s, and a final elongation stage at 72 °C for 5 min were used for LCO1490/HCO2198 primers amplified samples. Positive and negative control were used each set of PCR sample runs. Ultrapure water, which replaces DNA templates, serves as a negative control and extracted DNA of *A. maculatus* was used as positive control. On 1.5% agarose gels, PCR products were visualized and positive reactions were verified by a clear band aligned parallel to 650 bp of the Invitrogen 100 bp DNA ladder (Thermo Fisher Scientific, MA) fragment. All positive PCR products were labelled with the acronym DADB1, DADB2,…DADB71, (Dietary Analysis using DNA Barcodes) (n=71) alongside positive and negative control and were commercially sequenced bi-directionally with the sequencing primers M13F and M13R (Ivanova et al., 2007) and BigDye Terminator Cycle sequencing kit (Applied Biosystems) on an ABI 3730 capillary sequencer at Macrogen Inc. (South Korea).

### 2.3. DNA sequence analysis

Until at least every generated sequence was >600 bp in length, sequencing efforts were repeated. Forward and reverse sequences were trimmed using ChromasLite ver.2.1 to remove ambiguous and/or low quality bases and remnant primer from amplification or sequencing reactions. Sequences were compiled, and by translating DNA sequences into putative amino acid sequences in BioEdit ver. 7.9 (Hall, 1999) and aligned in Clustal X ver. 2.0.6 (Thompson, 1997), the gaps within the DNA sequences were tested. More than 600 bp length were all final contigs. The Barcode of Life Data Systems (BOLD) ID search engine (Ratnasingham and Hebert, 2007) and GenBank’s BLAST tool (Altschul et al., 1990) have been used as a reference libraries to identify the barcode sequences generated. Molecular Evolutionary Genetic analysis (MEGA) ver. 4.1 (Kumar et al., 2018) using Kimura 2 parameters (K2P) (Kimura, 1980) was used to construct neighbour-joining (NJ) tree (Saitou et al., 1987). Bootstrap test (100 replicates) was used to test the reliability of the branches (Felsenstein, 1985). For tree based identification, sequences of statistically significant (highest query coverage (q) or lowest error (e) values) references were extracted from GenBank. For better representation of tree based identification, the NJ tree was redrawn using the Interactive Tree Of Life (iTOL) (Letunic and Bork, 2019). The K2P model and the Tajima’s test for pair-wise distance and nucleotide diversity (Tajima, 1989) estimation were conducted in MEGA, respectively. The compositions of nucleotides and the diversity of haplotype were measured using the workbench tools delivered in BOLD. Less than 3% of the divergence between the unknown and the reference sequence was used to assign a specie level identity (Valdez-Moreno et al., 2012). That is, if the sequence matches reference GenBank sequence by at least 97%, the species identity has been accepted. To estimate the number of species and haplotypes present in the sample, we used the accumulation curve provisions provided in BOLD to visualise the taxonomic and sequence diversity. The also helps the user to monitor and compare the efficacy of sampling between groups or sites.

Sequence data generated in this study was released through GenBank and accessible via accession numbers JX676110-JX676180. Sequences along with their meta-data were also made available via BOLD (www.barcodeoflife.org) and could be accessed via project code DADB, and title; “Dietary analysis using DNA barcodes” or through http://dx.doi.org/10.5883/DS-DADB.

## 3. Results

### 3.1. Morphometric characterization of *A. maculatus*

This is the first study that explores *A. maculatus’s* gut content that occurr in Indian water using DNA barcodes. Out of 40 *A. maculatus* sampled, average length of 21.15 cm and biomass of 130.27 g was recorded (Table 1). The total length varied from 11 and 30 cm, and biomass ranged between 112 and 151 g. In pre-and post-monsoons sampling, maximum (30 cm) and minimum (10.3 cm) lengths were recorded, respectively.

**Table 1:**
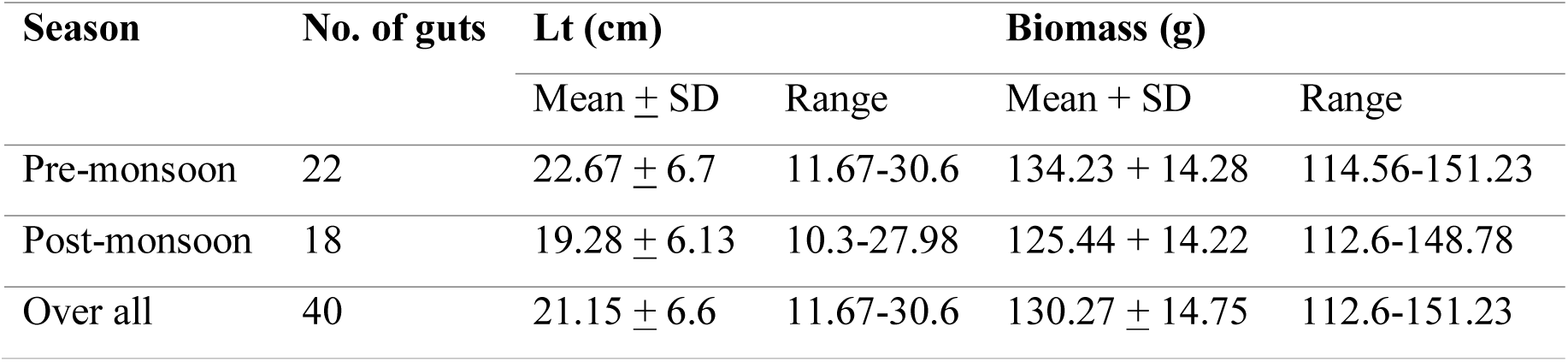
Mean values and ranges of *Arius maculatus* total length (Lt) and biomass sampled from the velar estuary

### 3.2. Dietary composition and sequence analysis

Of the 40 stomach contents examined (22 in pre-monsoon, 18 in post-monsoon), 32 stomachs (80%) had prey taxa that could be measured. 6 and 2 individuals collected in pre-monsoon and post-monsoon did not had measurable prey items, respectively. A total of 76 prey items were collected, of which 67 items were successfully sequenced (88.15%). The prey taxa could be classified into 4 major phyla, viz., Chordata, Arthropoda, Annelida and Mollusca. 4 barcodes of *A. maculatus* and 67 prey taxa constituted 71 sequences (DADB1 to DADB71) for BLAST analysis. DADB1 to DADB4 were *A. maculatus* tissue samples randomly sampled twice each time during pre-and post-monsoon. The pre-monsoon samplings of prey items were tagged from DADB5 to DADB33 (n=29) and the post­monsoon sampling were from DADB34 to DADB71 (n=38). **Table S1** list the respective PCR primer pairs (FishF1/FishR1 or LCO1490/HCO2198) used to barcode.

In GenBank database, all 71 sequences were matched with <3% cut off with that of reference sequences by identifying the taxa to species level. The prey taxa belonged to 13 species (12 genera in 11 families) under 4 phyla (viz., Chordate, Arthropod, Annelida and Mollusca) (The mean BLAST similarity score was 98.94). Pisces constituted the major prey items (43.28%) followed by Arthropod (29.85%), Annelida (17.91%) and Mollusca (8.95%). List of Pisces and invertebrate species and their seasonal variability was represented in Fig. 1. Of 29 Pisces sequences, 7 species were recognized *viz., Eubleekeria splendens, Stolephorus indicus, Photopectoralis bindus, Leptomelanosoma indicum, Lutjanus fulviflamma, Upeneus vittatus*, and *Gerres filamentosus*, respectively. In Annelida (*Capitella capitata, Perinereis vallata*), Arthropoda (*Clibanarius clibanarius, C. longitarsus*) and Mollusca (*Meretrix meretrix, Perna viridis*), the remaining taxa were made up of 2 species each.

**Fig. 1:**
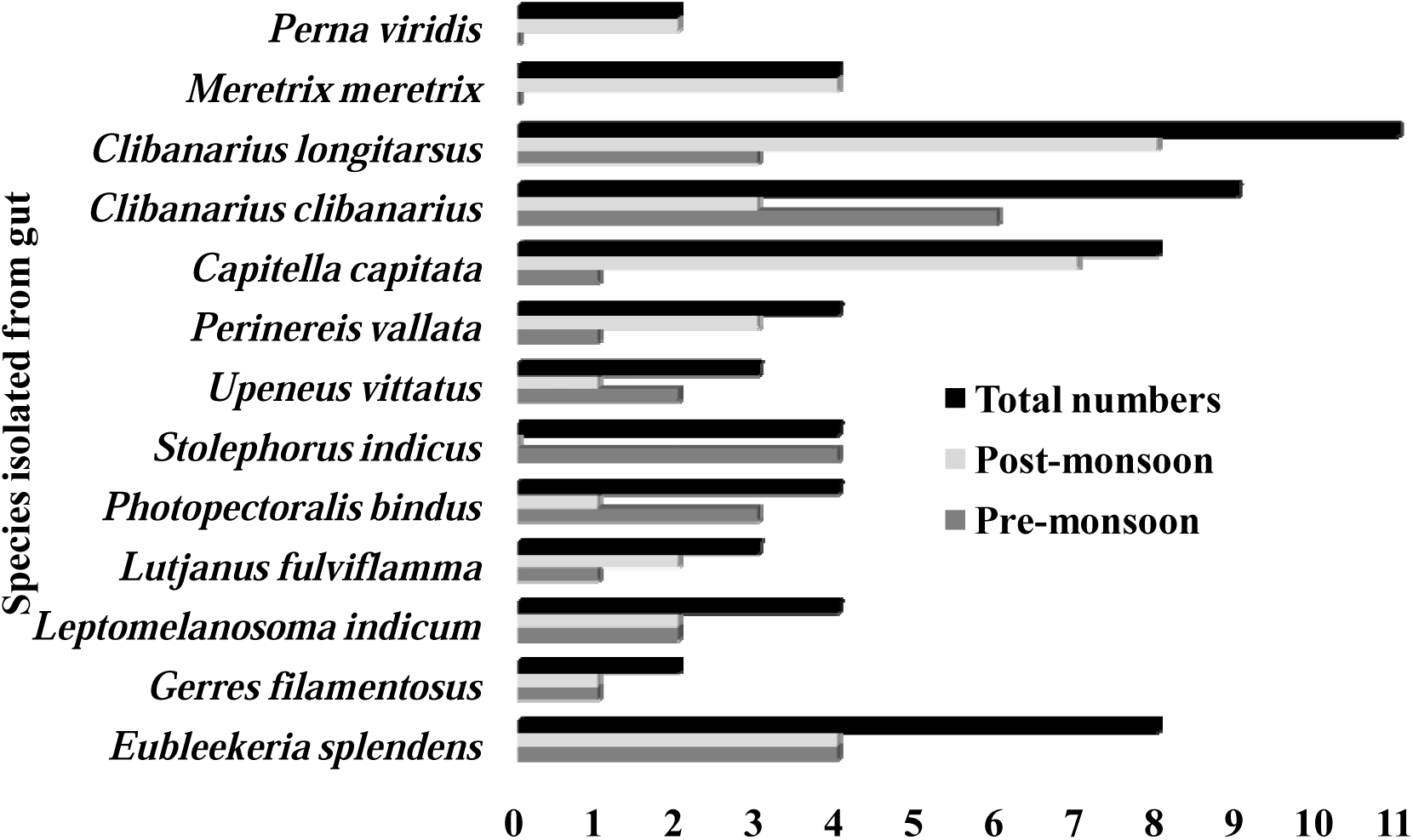
Seasonal variation in diet diversity of *A. maculatus.* The isolated species and their frequency of occurrence were given in the X and Y axes, respectively.

The mean BLAST similarity score was 98.94% with standard deviation of +0.99% (**Table S1**). The maximum and minimum similarity scores were respectively 100% and 97.01%. All 4 barcodes of *A. maculatus* was precisely identified with >99.3% identity (**Table S1**). All 71 barcodes produced in this study contained 43% average GC content (in the range of 35-50%) (Table 2).

**Table 2:** Summary statistics for nucleotide frequency distribution of dietary barcodes

| Composition (%) | Minimum | Mean | Maximum | Standard Error |
| --- | --- | --- | --- | --- |
| Guanine (G) | 16.98 | 19.12 | 22.73 | 0.22 |
| Cytosine (C) | 13.49 | 23.91 | 30.08 | 0.64 |
| Adenine (A) | 19.16 | 24.10 | 29.84 | 0.35 |
| Thymine (T) | 27.76 | 32.87 | 42.68 | 0.59 |
| GC | 35.04 | 43.03 | 50.86 | 0.56 |
| GC Codon position 1 | 40.47 | 52.15 | 59.45 | 0.64 |
| GC Codon position 2 | 38.14 | 42.63 | 44.08 | 0.13 |
| GC Codon position 3 | 19.52 | 34.31 | 52.58 | 1.08 |

### 3.3. NJ tree-based species identification

Identification of *A. macultus* specimens were also verified using the reference sequences extracted from GenBank via NJ tree construction. Other species in the *Arius* genera, such as *A. manillensis* (HQ682617), *A. dispar* (KF604635), A. *subrostratus* (MK348195), and *A. jella* (KU894613) was used as an out-group in the construction of NJ tree. All four sequences (DADB1-DADB4) have been placed in a single clade (Fig. 2). Presence of haplotypes within the 4 COI sequences of *A. maculatus* was also hinted.

**Fig. 2:**
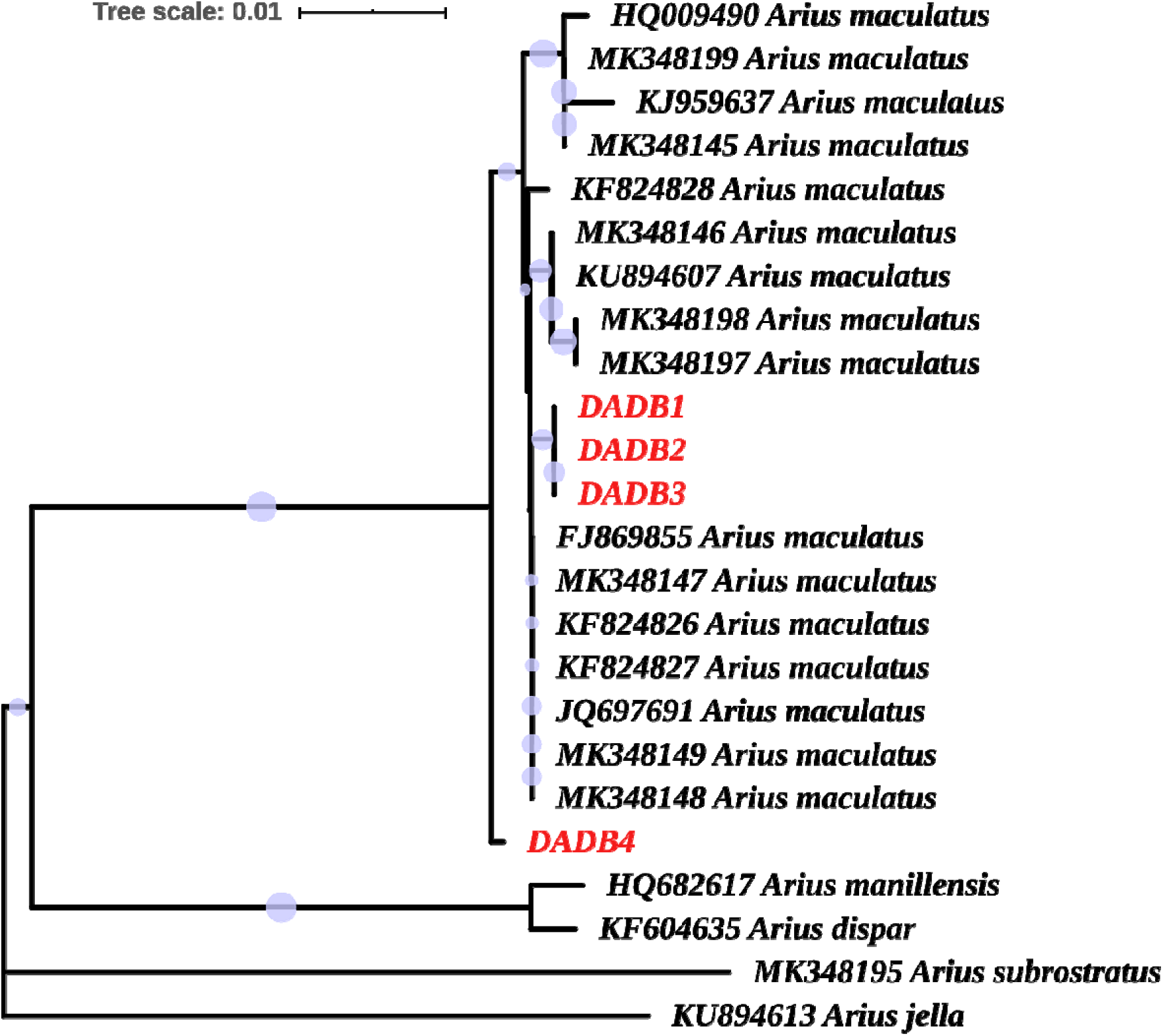
COI-NJ tree based identification of *Arius maculatus.* DADB1-DADB4 were the sequences produced in this study. The alpha-numerical present prefix to species name represent the Genbank accession numbers. Different species of *Arius* genera was used as an out-group was successfully delineated.

For the construction of the NJ tree, all 7 piscine species with their reference sequences (2 to 5 reference sequences per species) constituting a total of 56 nucleotide sequences were used. All pisces prey taxa clustered in one clade with its respective species indicating the similarity between the sequenced COI and sequenced taxa in the database (Fig. 3). Even the cladding patterns of constructed NJ tree suggested the presence of haplotypes among the prey taxa; for example, 2 sub-clades were evident among the *Stolephorus indicus* clade (the top most clade of the NJ tree) (Fig. 3). One contained DADB11, DADB12 and DADB24 with other sequences of reference, and another contained DADB26 and DADB30 with other sequences of reference. Similar patterns were also observed in *U. vittatus, E. splendens* and *L. indicum* clades. For the NJ tree based invertebrate prey taxa identification, the 2 to 9 reference sequence per invertebrate prey taxa was similarly used. The final dataset had 68 nucleotide sequences. All invertebrate prey taxa in Arthropod, Annelid and Mollusca, clustered with its respective sequences of the reference species in single clade (Fig. S1), indicating the success of tree based identification.

**Fig. 3:**
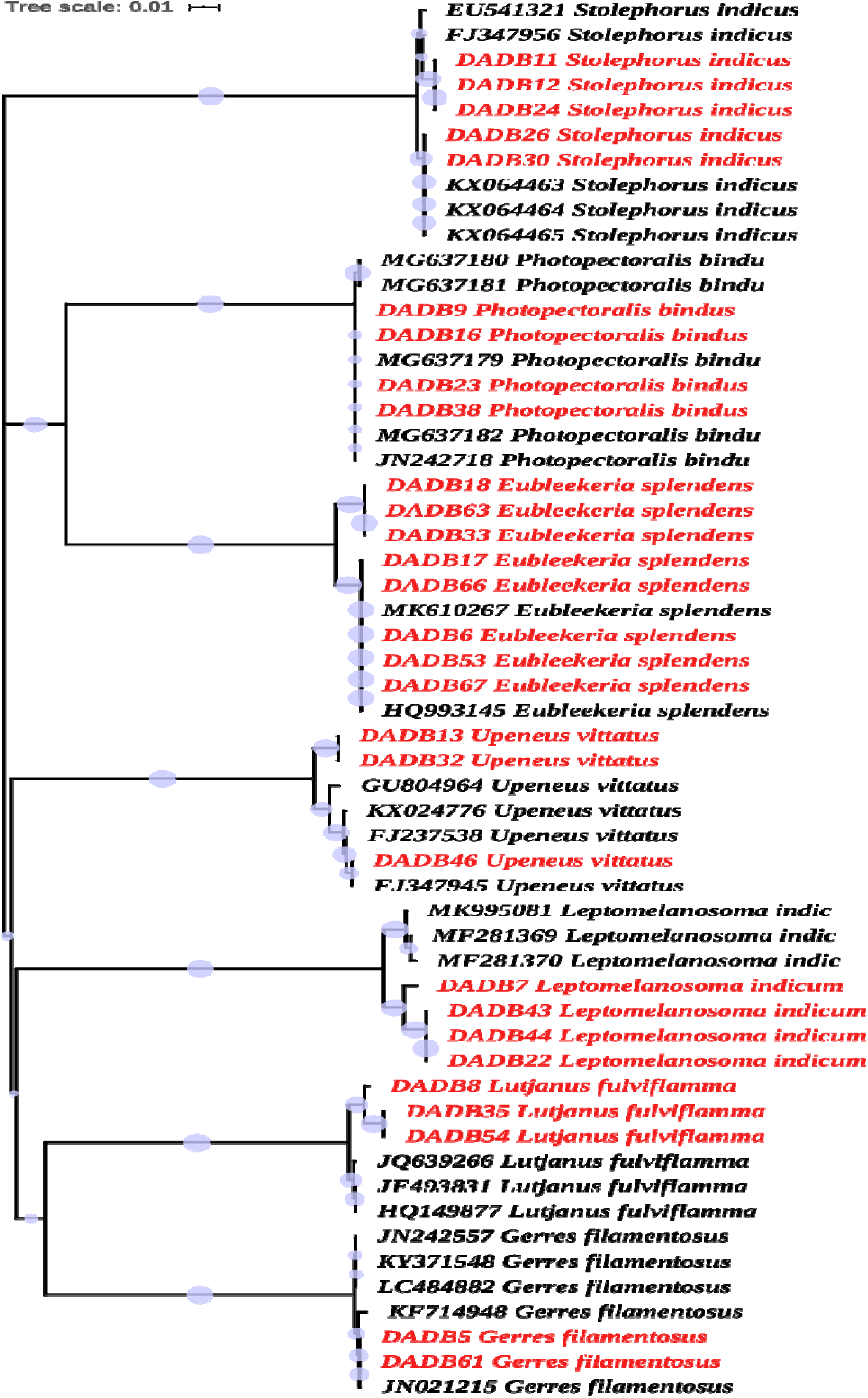
NJ tree constructed for tree based identification of pisces prey taxa. The percentage of replicate trees in which the associated taxa clustered together in the bootstrap test (100 replicates) are shown as circles next to the branches. The number of base substitutions per site was indicated as tree scale given on the top left corner. The acronym DADB and alpha-numerical prefix to the species name indicates the sequences produced in this study and reference sequence, respectively.

### 3.4. Dietary diversity: Pre-monsoon versus Post-monsoon

The major difference between pre-and post-monsoon dietary composition is that, in pre-monsoon, the molluscan taxa was completely absent and a piscine species *(Stolephorus indicus*) was present only in pre-monsoon (Fig. 1). Among piscine taxa, *Eubleekeria splendens* (23.5%) and *Stolephorus indicus* (23.5%) were the major prey taxa followed by *Photopectoralis bindus* (17.6%) during pre-monsoon while *E. splendens* (36.36%) alone forms the major prey taxa followed by *Leptomelanosoma indicum* (18.18%) and *Lutjanus fulviflamma* (18.18%) during the post-monsoon season.

The hermit crabs forms the major constituents of both pre-and post-monsoon seasons (81.81% & 40.74%, respectively) among invertebrate taxa. The dominant (54.54%) invertebrate prey taxa were the hermit crab, *Clibanarius clibanarius* was followed by *C*. *longitarsus* (27.27%) during pre-monsoon sampling. *Clibanarius longitarsus* were the dominant (33.33%) invertebrate prey taxa during post-monsoon sampling followed by a polychaete worm *Capitella capitata* (25.92%) (Fig. 1).

During pre-monsoon, piscine gut items constituted the major prey taxa (58.62%) and invertebrate forms the major prey (71.05%) taxa in post-monsoon season. During the post-monsoon, consumption of piscine items dropped by 20.69% and invertebrate taxa increased by 42.11%. However, as the sample size was severely limited to draw such inference, it was possible to adjust further sampling efforts accommodated with next generation sequencing to verify such claims.

For invertebrate taxa, the average pair-wise distance (pwd) and nucleotide diversity were higher (0.35 and 0.26, respectively) than for pisces (0.22 and 0.18, respectively) (Table 3). During the post-monsoon season, the total higher pwd values (0.40) were observed than pre-monsoon values (0.31) suggesting higher dietary diversity consumed in the post-monsoon season. Similarly, the decrease in pisces’s pwd and nucleotide diversity values and increased values of the same parameters in post-monsoon invertebrate prey taxa indicate that invertebrates were more preferred diet during the post-monsoon season. The overall pwd values for the barcoded prey taxa were 0.377.

**Table 3:**
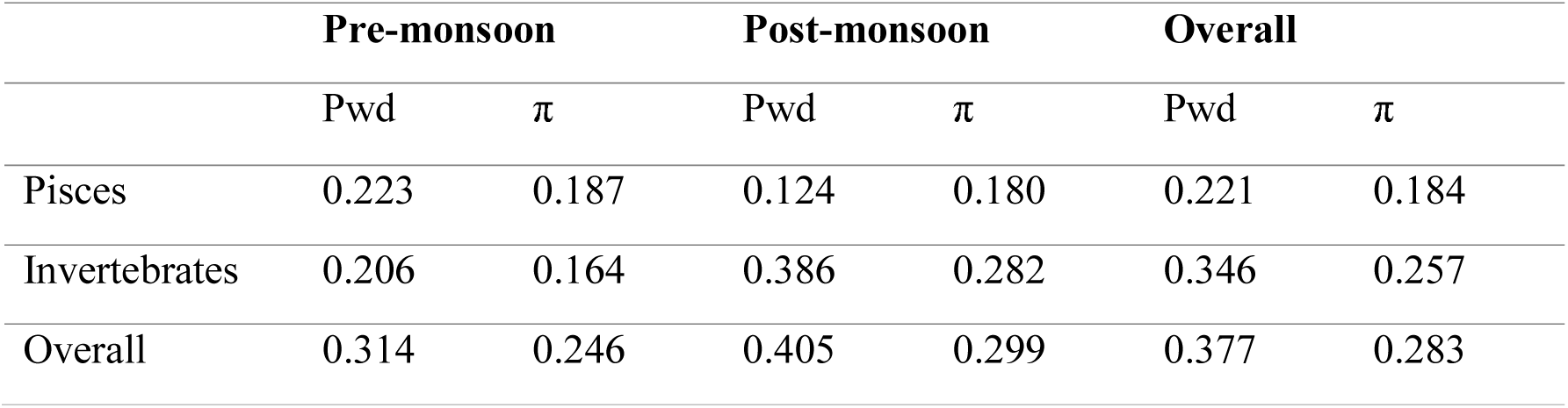
Pair-wise distance (Pwd) and nucleotide diversity (π) of diet-DNA barcodes

|  | Pre-monsoon |  | Post-monsoon |  | Overall |  |
| --- | --- | --- | --- | --- | --- | --- |
| | Pwd | $\pi$ | Pwd | $\pi$ | Pwd | $\pi$ |
| Pisces | 0.223 | 0.187 | 0.124 | 0.180 | 0.221 | 0.184 |
| Invertebrates | 0.206 | 0.164 | 0.386 | 0.282 | 0.346 | 0.257 |
| Overall | 0.314 | 0.246 | 0.405 | 0.299 | 0.377 | 0.283 |

### 3.5. Haplotype diversity

The analysis of the accumulation curve in BOLD reveals the presence of 57 haplotypes within 14 species barcoded (including the host *A. maculatus)* (Fig. 4). In Pisces (47.36%), the majority of haplotypes were found, followed by Arthropods (28.07%), Annelids (14.03%) and Mollusca (10.52%). Inthis analysis, the mean haplotype (n=57) richness per taxa barcoded taxa (n=14) was found to be 4.07.

**Fig. 4:**
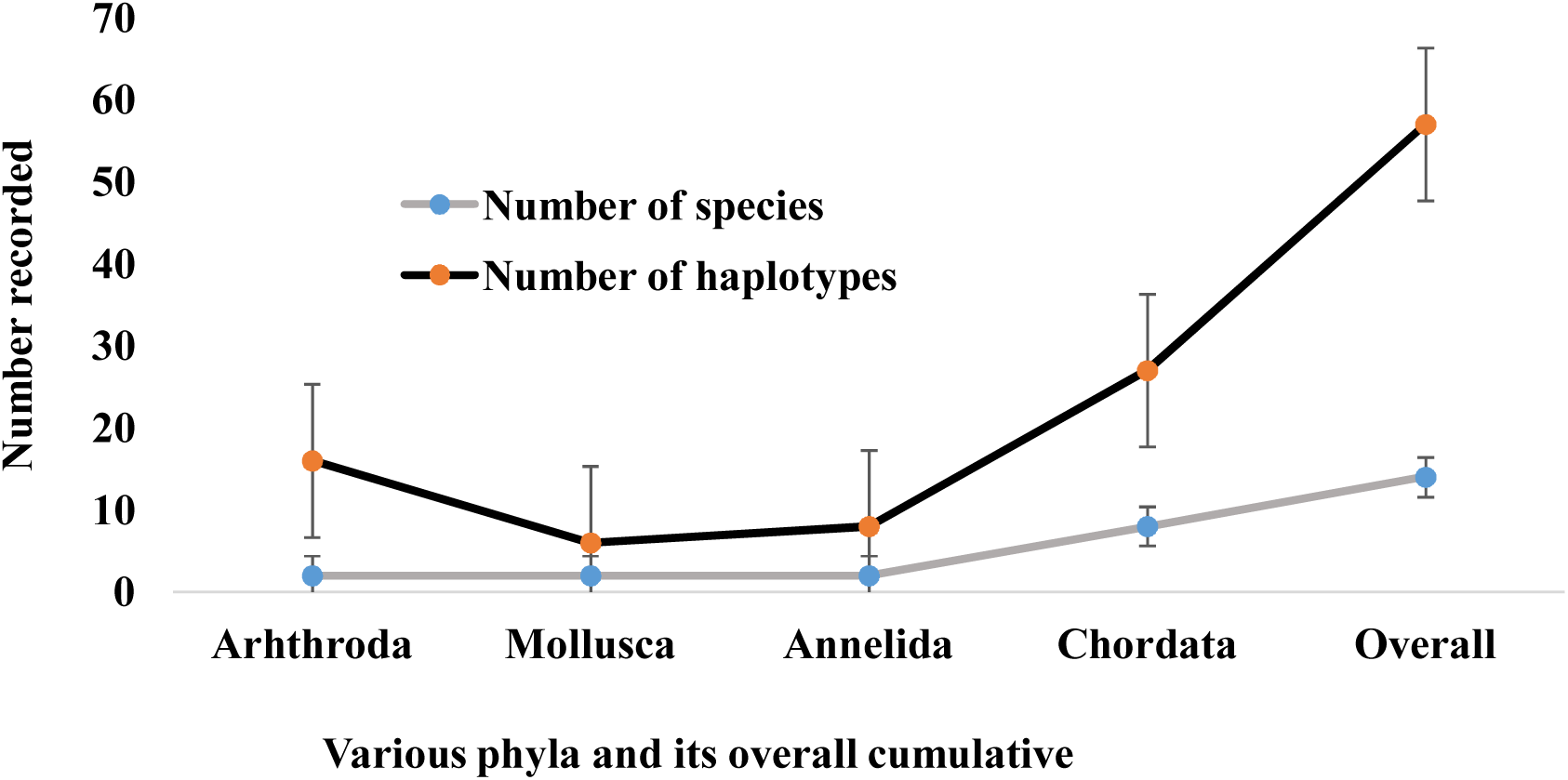
Accumulation curve demonstrating the documentation of various haplotypes present among the taxa in dietary composition.

## 4. Discussion and conclusion

In this study, the mean length recorded was comparable to *A. maculatus* occurring on the west coast of India (Maitra et al., 2019), but comparatively smaller than those occurring on the coast of Philippines (Chu et al., 2011). Many studies have not been able to identify the stomach content (prey items) of fishes using conventional morphological analysis (Legler et al. 2010; Paquin et al. 2014; Moran et al. 2015), as the digestive process rapidly disintegrates the morphological characteristics of the prey item (Schooley et al. 2008; Legler et al. 2010; Carreon-Martinez et al. 2011), resulting in substantial losses of the valuable data for taxonomist and resource managers. While in this study, most prey items were in high digestive state and rendered morphological identification impossible, segregating and treating the individual stomach content for DNA barcoding resulted in a success rate of 88% sequencing with 100% species level identification. Using visual survey, previous analysis, which could only assign 10% of catfish gut content to species level, used DNA barcoding to witness the taxonomic assignments to species level identification of 90% of prey items (Aguilar et al., 2016). None of the specimen could be recognised in this study without DNA barcoding, which could lead to poor understanding of catfish and their diet diversity. Previous nationalised efforts to barcoding the marine diversity (Lakra et al., 2010; Bineesh et al., 2014; Bamaniya et al., 2015) along with localised efforts to barcode the diversity of Vellar estuary (Khan et al., 2010, 2011; PrasannaKumar et al., 2012; Thirumaraiselvi et al., 2015; Rajthilak et al., 2015; Rahman et al., 2013, 2019; Hemalatha et al., 2016; Sahu et al., 2016; Palanisamy et al., 2020; PrasannaKumar et al., 2020; Manikantan et al., 2020; Thangaraj et al., 2020; Narra et al., 2020) resulted in strengthening the reference library which insured that no ambiguous sequences were present in this study (as all sequences were identified to species level). Identification success was likely due to the use of previously generated references sequences from Indian waters from morphologically verified species and published through reference databases (such as GenBank and BOLD). This is significant because the ability of DNA barcoding to identify unknown specimens might be impeded by miss-identification of reference specimen, cryptic diversity, sharing of haplotype, and lack of reference sequences (i.e., species yet not barcoded) in the database.

Previously, cryptic diversity has been recorded among catfish prey (April et al., 2011) and Indian waters have vast marine fish diversity (2443 species) (Gopi and Mishra, 2015) with several cryptic families (Bamaniya et al., 2015). The application of DNA barcoding in this study, identified 57 haplotypes in 14 species barcoded. When we previously barcoded Vellar estuarine fishes (43 species), for the first time 58% (n=25 species) were sequenced (Khan et al., 2011). These first time barcodes have been useful to identify *Lutjanus fulviflamma, Stolephorus indicus, Upeneus vittatus,* and *Eubleekeria splendens* in this study. In this study, the wide diversity of marine fish in catfish diets (7 species in 5 families) was not surprising, as previous studies have witnessed 23-25 fish taxa (up to 11 families) in catfish diets (Moron et al., 2015; Aguilar et al., 2016). Previous studies using DNA barcoding to investigate the catfish diet were limited to fish prey items (Moron et al., 2015; Aguilar et al., 2016), considering the difficulties of segregating and sequencing invertebrates. This study is the first of its kind to investigate the diversity of invertebrate prey in the diet of catfish using DNA barcodes. The economically important clams *(Meritxix meritrix*) (Yeh et al., 2016; Desrita et al., 2019) and mussel *(Perna viridis)* (Sulvina et al., 2020) found in the diet of *A. maculatus* in this study is of concern, as local populations were known to depend on these resources for food. Since accurate identification of macro-benthic invertebrates from benthophagous fish could facilitate the assessment of human impact on the ecosystem (Tupinambás et al., 2015), further studies could be directed toward a detailed picture of trophic levels in DNA barcoding based gut analysis of other catfish and fin fish species inhabiting Vellar estuary (Khan et al., 2011; Sakthivel et al., 2012).

We found that in pre-monsoon *A. maculatus* was more piscivorous than post-monsoon. However such conclusions should be supported by multiple seasonal sampling and not with single seasonal sampling like in this study. It was also appropriate to note that most prey items (almost all larvae and immature forms) identified in this study, were of commercially important fish species such as *Eubleekeria splendens,* which occurred abundantly in the East and West coast of India (Rawat et al., 2019). The diversity of diet estimated through DNA barcoding shows that, *A. maculatus* can feed at a range of depths and habitats (especially indicated by the high haplotype diversity), including shallow margins (such as mangrove habitats where immature forms seeks habitats) and open waters. The DNA sequencing mechanism and success of the detection of prey items might be disrupted by co-amplificatin of the predator’s DNA along with its prey (Vestheim and Jarman 2008; Leray et al. 2015). We have not obtained any co-amplification of *A. maculatus,* however. Similar observations were also previously made (Carreon-Martinez et al., 2011). While contaminations may have triggered sequencing failures (since approximately 12% of the samples were not successfully sequenced), false positive detections did not result (as all prey barcodes matched the level of species cut off with that of the same species in the reference database).

We found that DNA barcoding was very effective in the identification of even highly digested prey items. In this and other studies, prey items have been digested to the point that even higher taxonomic ranks of the prey could not be given an indication by the visual exam (Carreon-Martinez et al. 2011; Schloesser et al. 2011; Moran et al. 2015; Rocha et al. 2015; Aguilar et al., 2016). Even though the present study involved the detection of fish and invertebrates through DNA barcoding, the success rates of barcoding were higher than previously reported ∼65% by Moran et al. (2015), 70% by Cote et al. (2013), ∼80% by Carreon-Martinez et al. (2011) and nearly equivalent to (93%) Aguilar et al. (2016). The variations in the success rate of barcoding may be due to the choice of sequencing techniques, predator/prey within each study, prey item acquisition, predator/prey handling, and prey condition (digestion resistant features) (Macdonald, 1982; Buckland et al., 2017). Although the DNA barcoding techniques for fish and invertebrate using universal robust primers have been well developed (Ivanova et al. 2007; Weigt et al. 2012, Ward et al., 2005, Prasannakumar et al., 2012, 2020), barcoding success rates may be more likely to be affected during prey acquisition and other upstream barcoding processes (storage, pre-processing, DNA extraction, etc.), as higher success rate was witnessed in processing freshly acquired guts rather than the ethanol preserved whole predator samples. Limited diffusion of preservative medium into gut contents was reported for decrease success rate (Valdez-Moreno et al., 2012). In this study, care was taken to process the gut samples as fresh as could and the individual prey items was stored in ethanol rather than the whole predator. We also recommend that the extracted gut contents be stored in a deep freezing condition rather than in a preservative medium for a higher success rate of barcoding as previously observed (Aguilar et al., 2016).

We propose estimating the degree of generalist predation among the predators through pair-wise distance (pwd) and nucleotide diversity (π) estimation. For example, in post-monsoon in the diet of *A. maculatus,* the pwd and nucleotide values of prey items increases, indicating more post-monsoon generalist predation than pre-monsoon predation. However, it was beyond the scope of this study to investigate the factors responsible for such a rise. For now, the influence of pwd and π values on predator’s biology, functioning of predator habitats is unexplored, as more studies may be directed towards it in the near future. These barcode data (along with the associated full diet analysis) will reveal *A. maculatus’* trophic dynamics in Vellar estuary and provide valuable data for the development of management strategies, particularly in relation to its predation of commercially important fish and invertebrates. Pisces and invertebrates from these ecosystems will continue to be collected and barcoded, as the coverage of species barcodes in the reference database will become more significant as meta-and environmental DNA barcoding is becoming cheaper and more prevalent in fishery surveys (Leray et al. 2013; Miya et al. 2015; Galal-Khallaf et al., 2016; Evans & Lamberti, 2018).

## Acknowledgement

First author thanks the Department of Science and Technology’s INSPIRE Fellowship (IF10431), India and China Postdoctoral Research Foundation’s National Postdoctoral fellowship (0050-K83008), China for the financial assistance. The field help rendered my Mr. Karuna is acknowledged. Authors are grateful to anonymous reviewers for improving the quality of this manuscript.

